# Marine oil snow formation in microcosm incubations of artificial seawater at 4 °C

**DOI:** 10.64898/2026.11.02.763553

**Authors:** Thamali Vidanage, R. Eric Collins

## Abstract

An oil spill in the Arctic would have far-reaching impacts on the environment, the biota, and the Inuit communities that depend on this unique ecosystem for their livelihoods. Due to the rapid retreat of sea ice in the region, the Arctic is becoming more vulnerable to oil spills. Since microorganisms are the first responders to oil contamination, understanding their behavior is critical for designing bioremediation approaches tailored to Arctic conditions. In this study, we conducted microcosm experiments at 4 °C in the dark to simulate an Arctic oil spill, using crude oil and diesel as hydrocarbon sources. Biostimulation was tested with the addition of nutrients NaNO_3_ and KH_2_PO_4_, and soy lecithin–Tween 80 dissolved in ethanol as an alternative dispersant. Microcosms were placed on tube rollers at 3 rpm to mimic natural wave action and run with appropriate controls to assess microbial community shifts. Our objectives were to characterize the early microbial succession associated with oil degradation and marine oil snow (MOS) formation, and to determine how nutrients, dispersants, oil type (crude vs. diesel), photochemical aging (fresh vs. weathered), and oil concentration influence these processes.

We observed that the combined effect of nutrients and dispersant resulted in rapid MOS formation, occurring as early as 30 minutes after the initial setup. Microbial taxa exhibited varying tolerance to different concentrations of the same oil, and the dominant hydrocarbon degraders in fresh diesel treatments were distinctly different from those in weathered diesel. Furthermore, the nutrient–dispersant treatment promoted an earlier succession of oil-degrading taxa compared to nutrient-only or dispersant- only treatments. Together, these results highlight the importance of cold-adapted microbial communities in driving early oil degradation and MOS formation in Arctic waters, and they provide insights relevant to the development of Arctic-specific bioremediation strategies.

## 1 Introduction

The Arctic is at risk from future oil spills due to increased vessel traffic elevated by the rapid retreat of ice cover (Kapsar et al., 2023). Northern communities rely heavily upon ships as a way of transporting goods to serve the region (Prowse et al., 2009; Pelletier and Guy, 2015), but the continued expansion of maritime activities in the Arctic is anticipated to raise the risk of oil spills (Afenyo et al., 2019), which pose acute and chronic stressors to the environment, wildlife, and Inuit communities living there (Wright et al., 2022a). The composition of spilled oil changes immediately after its release into water (Mariano et al., 2008). While spilled oil undergoes evaporation, photo-oxidation, and dispersion (Garrett et al., 1998), biological degradation is the primary process for removing oil from both surface and subsurface plumes (Scheibye et al., 2017). Breaking down hydrocarbons requires an appropriate consortium of degraders and conditions that support the growth and multiplication of these microorganisms (Ron and Rosenberg, 2014). Bioremediation in cold marine environments is considered a labor-effective technique since it does not produce harmful byproducts (Brakstad et al., 2017). However, the rate and extent of oil biodegradation are influenced by the availability of nitrogen (N) and phosphorus (P) in the marine environment (Hazen et al., 2016). Surface seawater typically contains low (nano- or micromolar) concentrations of nitrogen and phosphorus, and hydrocarbons themselves are deficient in these essential nutrients (Atlas and Bartha, 1973). Therefore, the addition of N and P following an oil spill has been shown to support the native marine microbiota that contributes to oil biodegradation (Mirsaeidi, 2023). Furthermore, the rate of biodegradation depends on various factors such as temperature, oil composition, presence of electron acceptors, and an active microbial population (Rahsepar et al., 2016a). Nevertheless, biodegradation rates are often overestimated (Mariano et al., 2008), particularly due to oil being photo-oxidized and volatilized, and by ocean currents moving the oil to shore (Ron and Rosenberg, 2014).

Since different oils have distinct physical, chemical, and toxicological characteristics, they behave differently in the marine environment and can produce markedly different environmental impacts when spilled (Wright et al., 2022b). After an oil spill, skimmers are typically used to remove oil mechanically. However, harsh weather conditions can severely limit their effectiveness, and even the largest booms and skimmers may be unable to operate, e.g., in harsh Arctic conditions (Prince et al., 2017). Hence, responders use dispersants to reduce the impact of large oil slicks reaching shorelines and to promote biodegradation (Prince, 2015). Chemical dispersants consist of surfactants and solvents (Wise and Wise, 2011) that reduce the interfacial tension between oil and seawater, forming smaller oil droplets and making oil highly accessible for microbial degradation (Lessard and DeMarco, 2000; Rahsepar et al., 2016a). Though the toxicity of dispersants varies depending on the dispersants and the species affected, oil concentration and type of oil dispersants may not pose a significant risk to aquatic species in field conditions (Bejarano, 2018). With advances in formulation chemistry, modern dispersants are far less toxic than earlier products (Kleindienst et al., 2015a), with food-grade and biological surfactants being promising eco-friendly dispersants, given their non-toxicity and biodegradability (Athas et al., 2014). Lecithin and Tween 80 are the most widely studied surfactants used as dispersants, owing to their enhanced emulsifying capacity (Jin et al., 2019). In this study, we used a lecithin–Tween 80 mixture at a 60:40 (w/w) ratio, dissolved in ethanol, as the dispersant. Lecithin is a mixture of phospholipids, glycolipids, triglycerides, and fatty acids, and Tween 80 (Polysorbate 80; C_64_H_124_O_26_) is a complex nonionic surfactant and emulsifier; both are used widely in food production and industrial applications (Riehm et al., 2017). Dispersants promote the solubility of spilled oil by increasing the surface-to-volume ratio (Prince, 2015), making them accessible for microbes (Kleindienst et al., 2015); aggregation can also affect oil–microbe interactions, e.g., the formation of marine oil snow (MOS), a key process influencing the aggregation, sedimentation, and microbial degradation of oil in the water column (Passow, 2016a; Quigg et al., 2020; Gregson et al., 2021).

This microcosm study was designed as a pilot project to explore the impacts of fresh and weathered diesel and crude oil on MOS-associated biodegradation under controlled Arctic-like conditions. Experimental treatments incorporated varying hydrocarbon concentrations, nutrient amendments, and dispersant regimes to simulate realistic spill scenarios. We aimed to characterize the early microbial responders to oil contamination and evaluate their potential functional roles in degrading different hydrocarbon sources. These microbial dynamics provide insight into the mechanisms governing MOS formation and its role in oil aggregation, vertical transport, and biodegradation in cold marine environments.

## 2 Materials and Methods

### 2.1 Field sampling

Artificial seawater samples were collected into a large sterile cubitainer from the Sea-Ice Environment Research Facility (SERF) at the University of Manitoba on April 7^th^, 2025. The SERF pool (60 ft *×* 30 ft *×* 8 ft) had been filled with an artificial seawater mixture composed of a commercial brine slurry dissolved in groundwater to mimic seawater salt concentrations. The water sample was immediately stored at 4 °C in the cold room facility at the Centre for Earth Observation Science for 2 days until the microcosms were set up.

### 2.2 Microcosm setup and subsampling

The microcosms were established in 125 mL Pyrex glass bottles placed on tube rollers, following the method described by Gutierrez et al. (2013). Three sets of experimental groups were created with one replicate per sample, and each was monitored until the first signs of MOS were observed. The hydrocarbons used for this experiment included fresh Churchill diesel, UV-weathered Churchill diesel, and UV-weathered Tundra crude oil. Two concentrations were tested: 1% (v/v) and 0.1% (v/v).

Before the experiment, aliquots of Churchill diesel fuel and Tundra crude oil were exposed to UV weathering using UV-A (365 nm) and UV-B (302 nm) lamps for 5 days. A total of 24 microcosms were prepared, with each consisting of 98.9 mL of seawater amended with either UV-weathered Churchill diesel, UV-weathered crude oil, or fresh diesel at concentrations of 1% (v/v) or 0.1% (v/v).

The experimental setup included three treatment types for each oil and concentration: (1) nutrients only, (2) dispersants only, and (3) a combination of nutrients and dispersants. For dispersants, soy lecithin and Tween 80 (60/40, w/w) dissolved in ethanol were added at a final concentration of 0.1% (v/v) to the appropriate treatments. Two control treatments containing only seawater and UV-weathered diesel were prepared at 0.1% (v/v) and 1% (v/v).

Microcosms that contained nutrients were amended with 7.785 mM NaNO_3_ and 0.2 mM KH_2_PO_4_ to provide nutrient-replete conditions (1000*×* the natural environment) and satisfy nutrient requirements based on Redfield ratios. Microcosms were placed horizontally on tube rollers set at 3 rpm and maintained at 4 °C in the dark to simulate natural oceanic conditions. All treatments, except those amended with fresh diesel, were destructively sampled after one week, coinciding with the formation of a substantial amount of MOS in the majority of the treatments. Fresh diesel-amended treatments were sampled after 3 weeks, once a comparable amount of MOS had developed. Observations were recorded at the time of sampling.

### 2.3 Genomic DNA extraction

Since the majority of MOS were at an early stage of development, they were not able to be separated reliably. Hence, the whole water sample was filtered onto a 0.22 µm Sterivex filter for genomic DNA extraction. The Qiagen DNeasy PowerSoil kit was used to extract DNA from the samples following the manufacturer’s instructions, with slight modifications. For all extractions, the bead-beating tube containing the samples and CD1 solution were heated to 70 °C for 10 minutes to facilitate cell lysis. Additionally, an aliquot of elution buffer was heated to 50 °C before eluting the DNA in the final step. All extracted DNA products were quantified using a Qubit fluorometer and stored at *−*70 °C until library preparation and sequencing.

### 2.4 16S rRNA amplicon sequencing

The V4 hypervariable region of the 16S rRNA gene was amplified using the 515F (GTGYCAGCMGCCGCGGTAA) and 806RB (GGACTACNVGGGTWTCTAAT) primer pair (CD Genomics, New York, USA). Amplicon libraries were sequenced using paired-end (2 *×* 250 bp) Illumina NovaSeq (CD Genomics, New York, USA). Approximately 680k high-quality reads were obtained with a read depth ranging from 14182 to 45158 per sample. Sequencing reads were first quality-filtered to remove adapters and primer sequences using Cutadapt v5.2 (Martin, 2011). The resulting trimmed reads were processed with the DADA2 package in R (Callahan et al., 2016) to identify amplicon sequence variants (ASVs), followed by taxonomic classification using the SILVA v138.2 reference database (Quast et al., 2013). Code for the bioinformatics pipeline is available at https://github.com/rec3141/microscape.

### 2.5 Statistical analysis and visualization

Microbial community composition analysis was performed using R. All plots were generated in R (v.4.3.1) using the ggplot2 package (Wickham, 2016).

## 3 Results

### 3.1 MOS development

The earliest development of MOS in roller incubations occurred within 30 minutes in the treatment containing 1% (v/v) diesel fuel, supplemented with 0.1% (v/v) dispersants and nutrients (Figure 1). In the weathered diesel treatments, MOS particles were suspended in the water column and were off-white in color, and were approximately 2 to 3 mm in size. However, after one week, they aggregated and became negatively buoyant and had settled at the bottom. The MOS produced in fresh diesel treatments differed significantly from that formed in weathered diesel treatments. The latter were noticeably whitish and had a mucous texture, remaining suspended throughout the water column, and small oil droplets were observed on the surface of the water. The shapes of those two categories varied, with some being linear and some being round. In weathered crude oil treatments, MOS particles were light brown in color, round in shape, and had negative buoyancy. The samples also contained linear structures that appeared to be an early stage of MOS development, which were very neutrally buoyant and suspended at the surface (Table 1).

**Figure 1.**
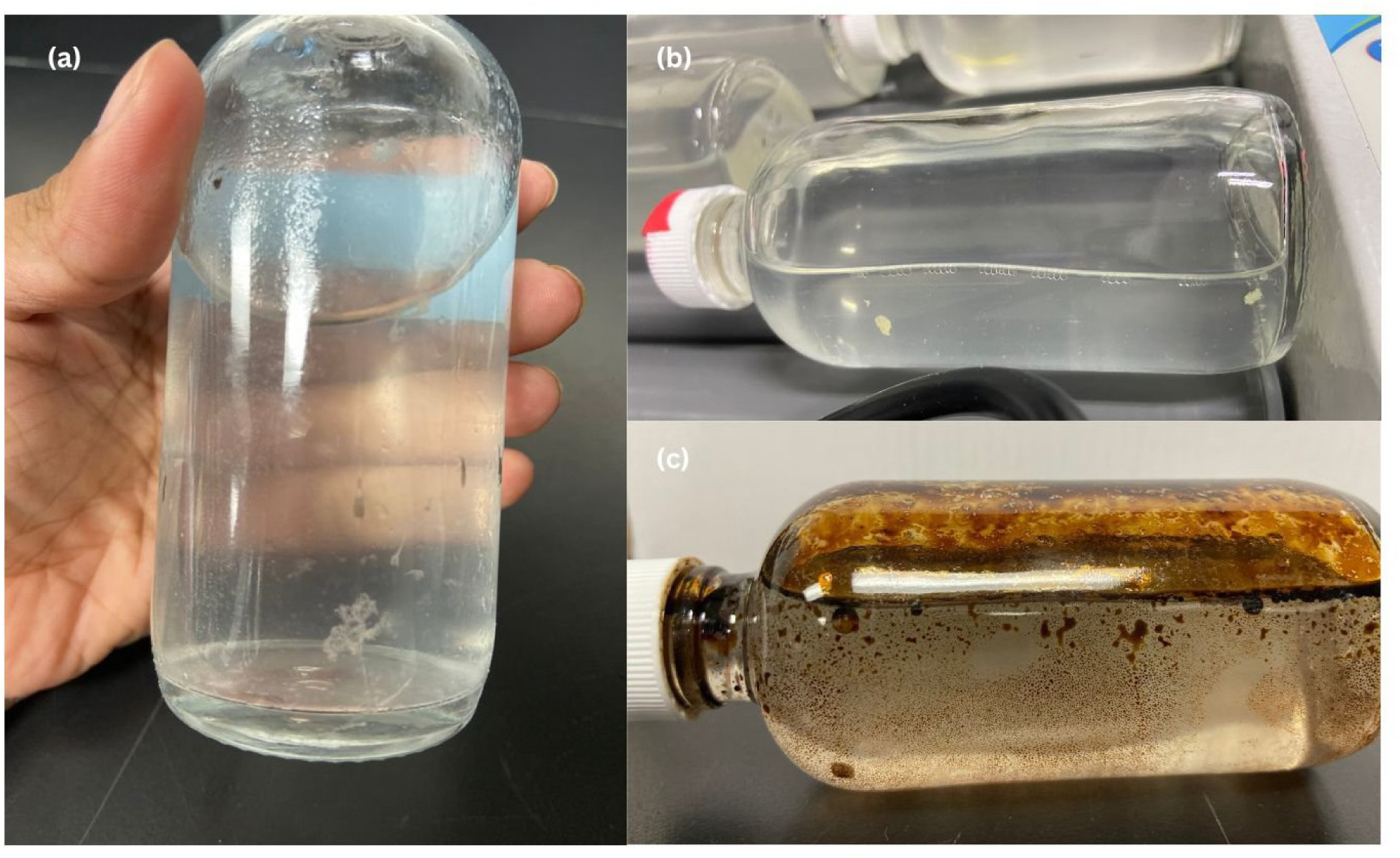
MOS aggregates formed in (a) fresh diesel (3 weeks old), (b) UV-weathered diesel (1 week old), and (c) UV-weathered crude oil treatments (1 week old).

**Table 1.** Experimental design for microcosm setup and visual observations of MOS formation

| Time | Treatment Group | Oil type | Oil: water ratio (%) | MOS presence | Observations |
|---|---|---|---|---|---|
| 3 WEEKS | SW + Oil + N | Fresh Diesel | 0.1 | Y | Tiny MOS particles scattered in the water column |
|  | SW + Oil + D | Fresh Diesel | 0.1 | N | Cloudy water, small oil droplets on the surface water |
|  | SW + Oil + N + D | Fresh Diesel | 0.1 | Y | Mesh-textured MOS, neutral buoyancy |
|  | SW + Oil + N | Fresh Diesel | 1 | N | Oil droplets on the surface water |
|  | SW + Oil + D | Fresh Diesel | 1 | N | - |
|  | SW + Oil + N + D | Fresh Diesel | 1 | N | - |
| 1 WEEK | SW + Oil + N | Weathered Diesel | 0.1 | N | - |
|  | SW + Oil + D | Weathered Diesel | 0.1 | N | - |
|  | SW + Oil + N + D | Weathered Diesel | 0.1 | N | - |
|  | SW + Oil + N | Weathered Diesel | 1 | N | - |
|  | SW + Oil | Weathered Diesel | 0.1 | N | - |
|  | SW + Oil + D | Weathered Diesel | 1 | N | - |
|  | SW + Oil + N + D | Weathered Diesel | 1 | Y | Off-white and dense MOS sunk in the bottom |
| 1 WEEK | SW + Oil + N | Weathered Crude Oil | 0.1 | Y | Linear structured MOS, positively buoyant. |
|  | SW + Oil + D | Weathered Crude Oil | 0.1 | N | - |
|  | SW + Oil + N + D | Weathered Crude Oil | 0.1 | Y | A light brown MOS was suspended for a few minutes |
|  | SW + Oil + N | Weathered Crude Oil | 1 | Y | Round-shaped MOS, positively buoyant |
|  | SW + Oil + D | Weathered Crude Oil | 1 | N | - |
|  | SW + Oil + N + D | Weathered Crude Oil | 1 | Y | A light brown MOS, negatively buoyant |

### 3.2 Bacterial community response to hydrocarbon inputs and biostimulation

Sequences clustered at 100% similarity yielded 243 ASVs, representing 10 phyla, 13 classes, 44 orders, 74 families, and 100 genera. Microbial community composition shifted significantly from the unincubated seawater (pre-treatment) to the amended treatments, indicating strong selective effects of oil exposure and experimental manipulations (Figure 2).

**Figure 2.**
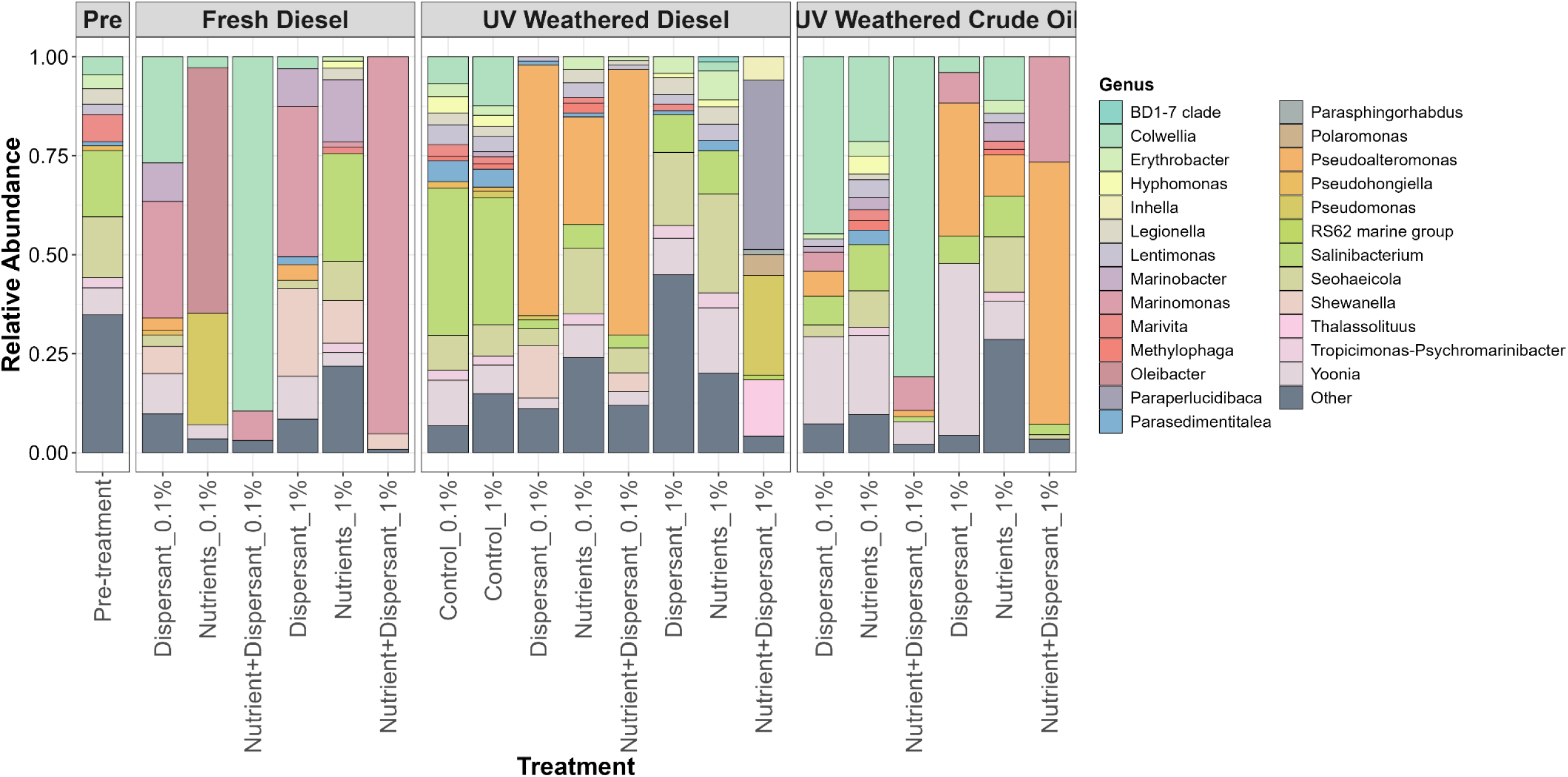
Bacterial community composition at the genus level for the unincubated seawater (pre-treatment), fresh diesel (after 3 weeks of incubation), control, UV-weathered diesel, and UV-weathered crude oil treatments (after 1 week of incubation). The hydrocarbon concentrations are shown as percentages (0.1% or 1%) with each treatment type. Each treatment had only one sample (n=1). shift in taxonomic dominance under UV-weathered oil exposure.

Control treatments containing UV-weathered diesel at 0.1% (v/v) and 1% (v/v) displayed nearly identical community structures at the time of sampling. In both controls, *Salinibacterium* dominated the communities (37.2% and 32.1%, respectively), suggesting limited compositional differentiation in the absence of nutrient and dispersant amendments.

In contrast, the addition of nutrients and dispersants induced pronounced shifts in community structure, indicating strong selective enrichment of hydrocarbon-degrading taxa. Treatments amended with fresh diesel at 0.1% and 1% (v/v) were dominated by distinct microbial taxa, reflecting concentration-dependent ecological selection. The 0.1% treatment was overwhelmingly dominated by *Colwellia* (89.5%), whereas the 1% treatment was dominated by *Marinomonas* (95.2%), suggesting that hydrocarbon concentration, in combination with nutrient and dispersant addition, exerted a powerful structuring effect on the microbial community.

Similarly, the nutrient-only treatments at 0.1% and 1% (v/v) differed markedly in community composition, suggesting a concentration-dependent response. The 1% treatment was dominated by *Oleibacter* (62.0%), suggesting strong enrichment of hydrocarbon-associated taxa under higher substrate availability. In contrast, the 0.1% treatment exhibited a more diverse community structure, with *Salinibacterium* (27.3%) as the most abundant taxon, followed by *Marinobacter* (15.6%), *Shewanella* (10.8%), and *Seohaeicola* (9.9%). These differences highlight the influence of hydrocarbon concentration on microbial selection even in the absence of dispersant addition.

Dispersant-only treatments exhibited broadly similar microbial community structures across the two diesel concentrations, although differences in relative abundances were evident. At 1% (v/v), *Marinomonas* was the most abundant taxon (38.0%), followed by *Shewanella* (22.2%), indicating strong enrichment of known copiotrophs at higher oil concentrations. In the 0.1% (v/v) treatment, *Marinomonas* remained highly abundant (29.4%), but at a level comparable to *Colwellia* (26.8%), suggesting a more balanced community structure at the lower diesel concentration. Overall, these results suggest that while dispersant addition shaped community composition consistently across concentrations, the hydrocarbon level also influenced the relative dominance of key taxa.

Similar compositional divergences were observed in the UV-weathered diesel treatments. At the lower weath- ered diesel concentration (0.1% v/v), treatments amended with dispersants alone and with the combination of nutrients and dispersants were dominated by *Pseudoalteromonas*, with relative abundances of 63.3% and 67.1%, respectively. In contrast, the corresponding treatments at higher diesel concentration (1% v/v) exhibited distinct community structures. Notably, *Paraperlucidibaca* was most abundant in the nutrient- and dispersant-amended treatment at 1% (v/v) weathered diesel (42.8%), suggesting a concentration-dependent

Clear divergence was also observed between fresh and UV-weathered diesel treatments, highlighting the influence of oil type on microbial assembly (Figure 3). Weathered crude oil treatments after one week exhibited distinct microbial community compositions compared to the pre-treatment (unincubated seawater), suggesting rapid shifts in response to crude oil exposure. Treatments amended with nutrients and dispersants at 0.1% and 1% (v/v) weathered crude oil supported markedly different microbial assemblages, suggesting concentration-dependent selection. At the lower crude oil concentration (0.1% v/v), *Colwellia* dominated the community (80.8% relative abundance), whereas at the higher concentration (1% v/v), *Pseudoalteromonas* was most abundant (66.2%). These results suggest that increasing hydrocarbon concentration under nutrient- and dispersant-amended conditions strongly influenced taxonomic dominance and community structure.

**Figure 3.**
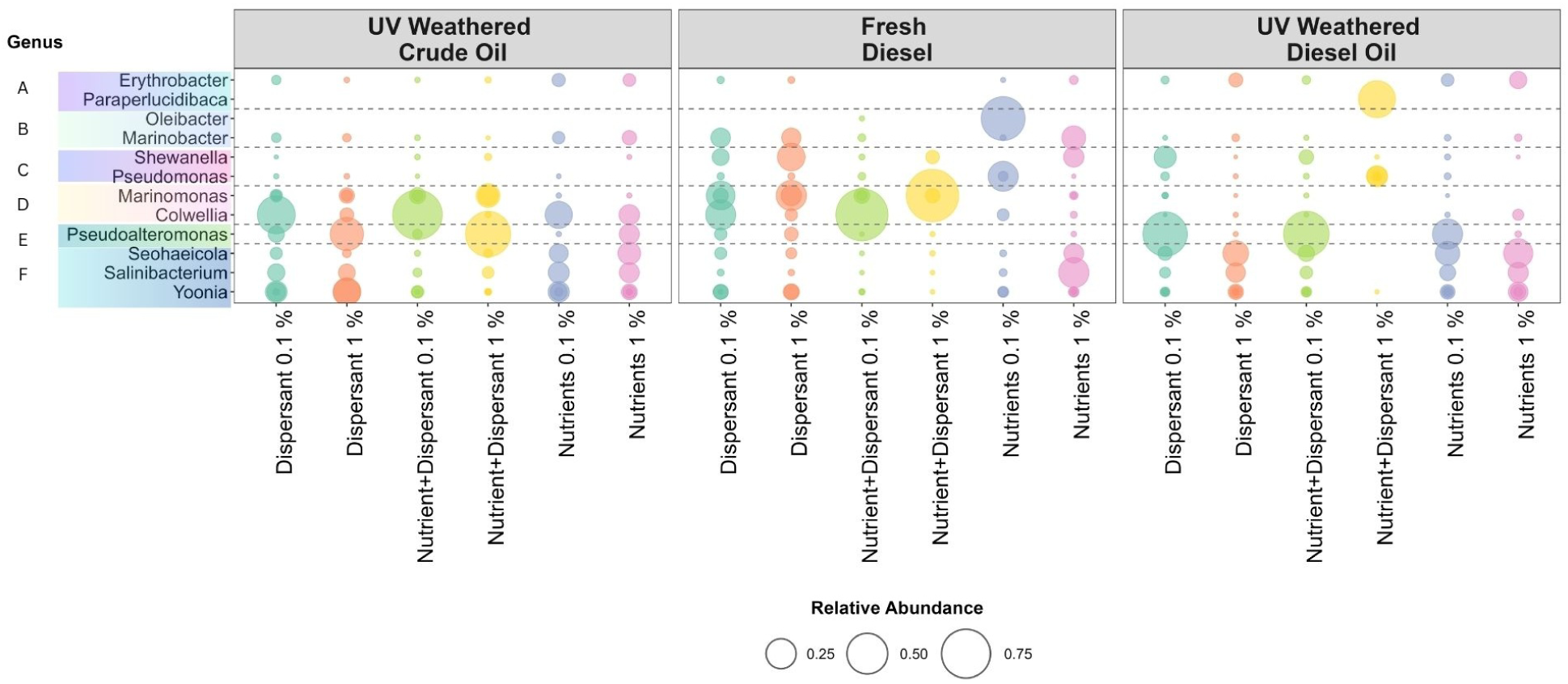
Relative abundance of bacterial taxa across different hydrocarbon inputs, concentrations, and biostimulation conditions. Bubble size is proportional to the relative abundance (%). Taxa were considered prominent if their relative abundance reached *≥*0.05% in any treatment and were grouped according to their distribution patterns: (A) prominent only in UV-weathered diesel, (B) in fresh diesel, (C) in both fresh and UV-weathered diesel, (D) in both UV-weathered crude oil and fresh diesel, (E) in both UV-weathered diesel and UV-weathered crude oil, and (F) in all three treatments.

Dispersant-only treatments exhibited greater variability in microbial community composition between the two crude oil concentrations. At the lower crude oil concentration (0.1% v/v), communities were characterized by the presence of *Colwellia*, *Yoonia*, *Salinibacterium*, and *Pseudoalteromonas*. In contrast, the higher crude oil concentration (1% v/v) was predominantly dominated by *Yoonia* and *Pseudoalteromonas*, indicating a concentration-dependent shift in community structure under dispersant exposure.

In contrast, nutrient-only treatments after one week at both crude oil concentrations supported more diverse microbial communities overall, while largely maintaining a community composition similar to that of the unincubated seawater. This suggests that nutrient addition alone exerted a relatively limited selective pressure compared to treatments amended with both nutrients and dispersants.

### 3.3 Bacterial community responses to hydrocarbon weathering and biostimulation

Specific bacterial taxa were associated with distinct hydrocarbon substrates, concentrations, and biostimula- tion regimes (Figure 3). In particular, *Marinobacter* and *Oleibacter* were detected in comparatively higher abundance in fresh diesel treatments, although their individual relative abundances varied across treatment types, suggesting differential affinity or tolerance to fresh diesel concentration and biostimulation conditions. Notably, *Oleibacter* exhibited the highest relative abundance in treatments containing the lower fresh diesel concentration amended with nutrients, indicating a potential preference for gaseous hydrocarbon availability under nutrient-enriched conditions.

In contrast, *Paraperlucidibaca* and *Erythrobacter* were detected at relatively higher abundances in weathered diesel treatments, suggesting substrate-specific selection or limited tolerance to fresh diesel and crude oil environments. *Pseudoalteromonas* dominated in weathered oil treatments (0.1% weathered diesel and 1% weathered crude oil), suggesting their preference for more complex or oxidized residuals and a later succession of this taxon in oil spills.

## 4 Discussion

Bacterial exudates play a significant role in the formation, modification, and degradation of marine oil snow (Simon et al., 2002; Grossart et al., 2003). Many microbes produce high molecular weight bacterial exudates in response to oil, which aid them in attachment, and also solubilize and emulsify the oil compounds, thus increasing the bioavailability of oil (Head et al., 2006a; McGenity et al., 2012). While the majority of the MOS studies have investigated the environmental effects, such as phytoplankton biomass, presence of particulate matter, and presence of EPS (extracellular polymeric substances) producing bacteria, not many studies have focused on how weathered oil with nutrients and dispersants would be involved in MOS formation.

The roller bottle incubations used in this study effectively simulated the formation of marine oil snow under varying hydrocarbon input concentrations and different biostimulatory conditions. Neither of the control treatments resulted in the formation of marine oil snow during the sampling period, whereas MOS formation was observed in 4/6 nutrient+dispersant treatments, 3/6 nutrient-only treatments, and 0/6 dispersant-only treatments.

In the present study, MOS formation was strongly influenced by the oil weathering state. Treatments containing fresh diesel required approximately three weeks for visible MOS formation, which formed in 2/6 treatments. In contrast, treatments with weathered diesel formed MOS more rarely (1/8 treatments), but they formed rapidly (within 30 minutes of incubation). Weathered crude oil produced MOS most frequently by far (4/6 treatments). These observations suggest that photochemically aged oil has an enhanced propensity to aggregate and form MOS. This finding is consistent with previous reports, including Passow (2016b), which demonstrated that photochemically aged oil promotes rapid MOS formation. The physical characteristics of MOS also differed markedly between treatments. Marine oil snow formed in weathered diesel treatments was denser and more compact, whereas MOS produced in fresh diesel treatments appeared loose and exhibited a more porous, mesh-like structure. These structural differences likely reflect alterations in oil chemistry following photo-oxidation, which may enhance aggregation potential and particle cohesion. Similar observations have been reported in microcosm experiments using photo-oxidized Macondo crude oil, where aggregates were smaller and more compact compared to those formed during the Deepwater Horizon oil spill (Passow, 2016b). Together, these findings suggest that weathering processes significantly influence not only the rate of MOS formation but also its physical structure and aggregation properties.

Though fundamentally limited in our interpretive power by the lack of replicates, the pronounced shifts in microbial community structure towards known oil-degrading taxa in nutrient and dispersant amendments do suggest a selective enrichment response to oil exposure. Similar restructuring of microbial assemblages after nutrient addition has been widely reported in oil-contaminated marine systems, where nitrogen and phosphorus limitation often constrain hydrocarbon biodegradation (Head et al., 2006b; McGenity et al., 2012). Nutrient enrichment can stimulate the rapid proliferation of obligate hydrocarbonoclastic bacteria, thereby accelerating biodegradation rates.

Dispersant addition has also been shown to alter microbial community composition by increasing hydrocarbon bioavailability through emulsification, although responses can vary depending on oil type and environmental conditions. For example, studies following the Deepwater Horizon oil spill documented enrichment of hydrocarbon-degrading taxa in response to dispersed oil fractions (Bælum et al., 2012). Our findings align with these observations, suggesting that the combined effect of nutrients and dispersants acts as a strong ecological filter, favoring taxa adapted to exploit dissolved and dispersed hydrocarbons.

Treatments containing the same hydrocarbon substrate but applied at different concentrations (0.1% and 1% v/v) under similar biostimulation regimes exhibited differences in microbial community composition. Without replicates, it is not possible to be certain whether these were stochastic batch effects or treatment effects, but the results suggest that oil concentration acted as a strong driver shaping microbial assemblages. Such concentration-dependent shifts likely reflect differences in bacterial tolerance thresholds, toxicity effects, and substrate availability. While lower concentrations may support a broader range of hydrocarbon-utilizing taxa, higher concentrations can impose physiological stress, selectively favoring taxa capable of tolerating elevated hydrocarbon exposure and associated toxic intermediates, as outlined in previous research (Zahed et al., 2010; Vilcáez et al., 2013).

The pronounced enrichment of *Colwellia* in fresh diesel treatments is consistent with previous observations that this genus rapidly responds to increased availability of gaseous and simple aromatic hydrocarbons (Mason et al., 2014). Fresh diesel is characterized by a relatively high proportion of volatile and low-molecular-weight aromatic compounds (Miracolo et al., 2010), which likely provided readily bioavailable substrates that favored *Colwellia* proliferation. The elevated abundance of *Colwellia* in treatments containing 0.1% (v/v) fresh diesel under nutrient and dispersant amendments further suggests that its growth was stimulated not only by substrate composition but also by enhanced hydrocarbon bioavailability and relieved nutrient limitation. Together, these findings support the view that *Colwellia* functions as an early responder to labile hydrocarbon inputs, particularly under conditions where dispersal and nutrient availability facilitate rapid substrate uptake.

*Marinomonas* exhibited significantly higher relative abundance in treatments containing 1% (v/v) fresh diesel, including those amended with nutrients and dispersants, as well as dispersant-only treatments, whereas nutrient-only amendments did not stimulate its growth. This pattern suggests that *Marinomonas* responds primarily to elevated hydrocarbon availability rather than nutrient enrichment alone. Supporting this observation, Zannotti et al. (2023) demonstrated that 1% (v/v) diesel can sustain *Marinomonas* growth, with optimal biodegradation performance reported at 22 °C. Although our incubations were conducted under colder conditions, the enrichment observed at higher diesel concentrations indicates that *Marinomonas* can tolerate and exploit substantial hydrocarbon loads even outside its reported temperature optimum.

Furthermore, consistent with our findings, Mugge et al. (2021) reported that *Marinomonas* proliferated in oil- and dispersant-treated surface seawater during a microcosm experiment evaluating biofilm responses to oil contamination. The enrichment of *Marinomonas* in dispersant-amended treatments in our study further suggests that increased hydrocarbon dispersion and bioavailability may facilitate its proliferation. Collectively, these results indicate that *Marinomonas* may be particularly adapted to environments with high hydrocarbon concentrations and enhanced oil dispersion, reinforcing its potential role as a concentration-responsive hydrocarbon-associated taxon.

The enrichment of *Pseudoalteromonas* in weathered and dispersant-amended treatments is consistent with previous reports describing its role in hydrocarbon biodegradation (Yakimov et al., 2004; Brakstad and Bonaunet, 2006). In addition to its metabolic capacity to degrade oil components, *Pseudoalteromonas* is known to produce EPS in cold marine environments (Suja et al., 2017). EPS production can enhance cell aggregation and provide a structural matrix that facilitates MOS formation (Passow et al., 2012). Therefore, its dominance in weathered oil treatments in the present study may not only reflect substrate utilization but also a functional contribution to aggregate formation and stabilization. This dual role in hydrocarbon degradation and EPS-mediated aggregation highlights *Pseudoalteromonas* as a potentially important taxon linking microbial succession with MOS dynamics under cold, oil-impacted conditions.

Comparison among treatment groups revealed that nutrient-amended treatments, irrespective of hydrocarbon concentration, exhibited microbial community compositions broadly similar to oil-only controls. This suggests that nutrient addition alone did not substantially restructure the microbial assemblage within the timeframe of this study. In oligotrophic environments such as the Arctic, hydrocarbon degradation is often limited by low nutrient availability; however, the response to nutrient enrichment may depend on the concurrent availability and physical state of hydrocarbons. Previous studies have shown that dispersant application can markedly enhance biodegradation by reducing oil droplet size and increasing the surface area-to-volume ratio, thereby improving microbial access to hydrocarbons (North et al., 2015). In contrast, nutrient addition without enhanced oil dispersion may not sufficiently alter substrate accessibility to drive pronounced community shifts.

Furthermore, significant restructuring of microbial communities under enhanced nutrient conditions during oil degradation in the Labrador Sea was reported to occur over approximately 28 days, often accompanied by accelerated hydrocarbon degradation (Murphy et al., 2021). In contrast, the present experiment was conducted over a one-week incubation period. It is therefore possible that this shorter timeframe was insufficient to capture more pronounced nutrient-driven community shifts, particularly in treatments where hydrocarbon bioavailability was not concurrently enhanced. These observations suggest that nutrient-mediated restructuring may require longer exposure periods to manifest fully under cold marine conditions.

Dispersant-only treatments, as well as combined dispersant and nutrient amendments, induced pronounced shifts in microbial community composition compared to oil-only controls within just one week, transitioning from a more generalist assemblage toward enrichment of specialized hydrocarbon-degrading taxa. This rapid restructuring suggests that dispersant-mediated increases in hydrocarbon bioavailability imposed strong selective pressure on the microbial community. The additional presence of nutrients may have further supported the proliferation of these specialized degraders by alleviating potential nutrient limitation. Together, these findings indicate that the combined application of dispersants and nutrients is likely to enhance the potential for hydrocarbon biodegradation within a short time.

The application of chemical dispersants as an oil remediation strategy remains controversial due to concerns regarding their potential long-term toxicity to marine fauna (Judson et al., 2010). Several studies have reported that dispersant addition can stimulate the growth of hydrocarbon-degrading bacteria (Dubinsky et al., 2013) and enhance overall biodegradation rates (Prince et al., 2016; Brakstad et al., 2018), primarily by increasing oil dispersion and bioavailability. In contrast, other investigations have suggested that dispersants may not enhance biodegradation and, in some cases, may inhibit the growth of certain hydrocarbon-degrading taxa (Lindstrom and Braddock, 2002; Kleindienst et al., 2015a; Rahsepar et al., 2016b).

In the present study, well-known hydrocarbon degraders thrived in dispersant-only treatments, even at elevated oil concentrations (1% v/v), suggesting that dispersant-mediated increases in substrate accessibility may have outweighed potential inhibitory effects under our experimental conditions. However, previous work has demonstrated that dispersants can differentially affect specific taxa. For example, Kleindienst et al. (2015b) reported that dispersant application altered competitive dynamics among hydrocarbon degraders, potentially suppressing taxa such as *Marinobacter* while favoring dispersant- and hydrocarbon-degrading bacteria, including *Colwellia*. Although our results show enrichment of several hydrocarbon-associated taxa under dispersant exposure, the present findings are unable to disentangle whether these shifts were driven by hydrocarbon utilization, dispersant degradation, or competitive interactions between taxa.

Furthermore, the ecological impact of dispersants is likely influenced by their chemical composition, as different formulations can variably shape microbial community assembly (Thomas et al., 2021). Collectively, these contrasting findings highlight the complexity of dispersant–microbe interactions and underscore the need to evaluate dispersant efficacy and ecological consequences under environmentally relevant conditions.

Although the present study did not directly quantify hydrocarbon biodegradation rates, the observed commu- nity restructuring implies that nutrient and dispersant amendments can substantially alter microbial assembly processes. Given these pronounced compositional shifts, the role of nutrients and dispersants may be critical in shaping microbial responses and ultimately influencing the environmental fate of spilled oil.

## 5 Conclusion

Overall, this study highlights the importance of combined nutrient and dispersant application as a potential bioremediation strategy for oil spills, particularly in facilitating the early succession of oil-degrading bacteria. The food-grade dispersant used in this study may also have contributed to the selective enrichment of certain bacterial genera, potentially conferring a competitive advantage to taxa capable of utilizing either hydrocarbons or dispersant components.

Furthermore, oil weathering appeared to influence bacterial community composition, likely due to alterations in hydrocarbon chemistry and the relative availability of different compound classes. Although several taxa were shared between weathered diesel and crude oil treatments, genera such as *Paraperlucidibaca*, *Thalassolituus*, *Pseudomonas*, and *Shewanella* were absent in crude oil treatments, indicating that oil type and composition significantly shape microbial community structure. These differences underscore the role of substrate characteristics, including physicochemical properties and weathering state, in driving microbial succession.

This study is limited by its microcosm design, which does not fully replicate natural environmental conditions, including continuous nutrient flux, dynamic gas exchange, and ongoing oil weathering and evaporation processes. Additionally, each treatment combination was represented by a single biological replicate, which restricts the ability to quantify within-treatment variability and limits the statistical applications for comparative analyses. Consequently, future research should investigate longer-term microbial succession under more environmentally realistic conditions and directly quantify biodegradation efficiencies across different stages of oil transformation to better evaluate the effectiveness of combined biostimulation strategies.

## Data availability

All raw 16S rRNA gene sequence reads generated from this experiment have been deposited at NCBI’s Sequence Read Archive (SRA) under the BioProject PRJNA1176269 (accession numbers SAMN54986490 to SAMN54986481).

## Author contributions

**Thamali Vidanage** – Conceptualization, Data curation, Formal analysis, Investigation, Methodology, Visual- ization, Writing – original draft, Writing – review & editing, Project administration

**R. Eric Collins** – Conceptualization, Methodology, Writing – review & editing, Validation, Supervision,

Funding acquisition, Project administration

## Acknowledgements

We would like to express our sincere gratitude to David Binne and Durell Desmond for their support in acquiring water samples from the Sea Ice Environment Research Facility (SERF) at the University of Manitoba. We are also grateful to Jocelyn Plouffe for the support in sample preparation.

